# Wide variabilities identified among spike proteins of SARS Cov2 globally-dominant variant identified

**DOI:** 10.1101/2020.09.26.314385

**Authors:** Abantika Pal, Aruna Pal

## Abstract

SARS Cov2 is a newly emerged virus causing pandemic with fatality and co-morbidity. The greatest limitations emerged is the lack of effective treatment and vaccination due to frequent mutations and reassortment of the virus, leading to evolvement of different strains. We identified a wide variability in the whole genome sequences as well as spike protein variants (responsible for binding with ACE2 receptor) of SARS Cov2 identified globally. Structural variations of spike proteins identified from representative countries from all the continents, seven of them have revealed genetically similar, may be regarded as the dominant type. Novel non-synonymous mutations as S247R, R408I, G612D, A930V and deletion detected at amino acid position 144. RMSD values ranging from 4.45 to 2.25 for the dominant variant spike1 with other spike proteins. This study is informative for future vaccine research and drug development with the dominant type.

## Introduction

The recent pandemics evolved with the characteristics of pneumonia, fever, difficult breathing, and ultimately renal failure or cardiac arrest causing death. The causal agent identified as SARS Cov2 virus. The outbreak was first reported in Wuhan province of China in December 2019, hence termed as Covid 19. WHO has declared the Novel Corona virus Disease (COVID-19) as a pandemic on 11^th^ March 2020 affecting 201 countries/territories/areas throughout the World, currently affecting 215 countries. The recent data as on current date at the end of June shows 484,925 number of deaths and affected population as 9,525,937 worldwide. However, the data also reveals 5,174,686 number of people have recovered (Worldometer, https://www.worldometers.info/coronavirus/). Apart from other human corona virus identified so far, the greatest concern with this recently evolving Covid 19 (SARS-Cov2) involves its high fatality and very high transmissibility and contagious nature.

Reports are available for seven coronaviruses for human. The RNA genome of SARS-CoV-2 has about 30,000 nucleotides, encoding for 29 proteins^**1**^. The structural proteins are the spike (S) protein, the envelope (E) protein, the membrane (M) protein, and the nucleocapsid (N) protein. There are other 16 non-structural proteins (NSP1-16), ORF3, ORF6, ORF7, ORF8 and ORF10. Three coronaviruses might have crossed species barriers from bat to civet cat (SARS-CoV) or camel (MERS-CoV) or pangolin (SARS-CoV-2), before crossing to human^**2**^, hence of great zoonotic importance.

The emerging pathogenic Coronaviruses were severe acute respiratory syndrome coronavirus (SARS-CoV), Middle East respiratory syndrome coronavirus (MERS-CoV), and very recently evolved SARS Cov2 (Covid 19). The first report for SARS-Cov infection was obtained from China in 2002, while MERS-CoV was evolved from Saudi Arabia in June 2012. However, the case fatality rate of SARS-CoV-2 (2-3%)^**3**^ was reported to be much lower compared to the SARS-CoV (11%)^**3**^ or MERS-CoV(34%)^**4**^. A comparative genome analysis for SARS Cov, MERS-Cov and SARS Cov 2 have been represented diagrammatically (Figure 1).

**Fig 1:**
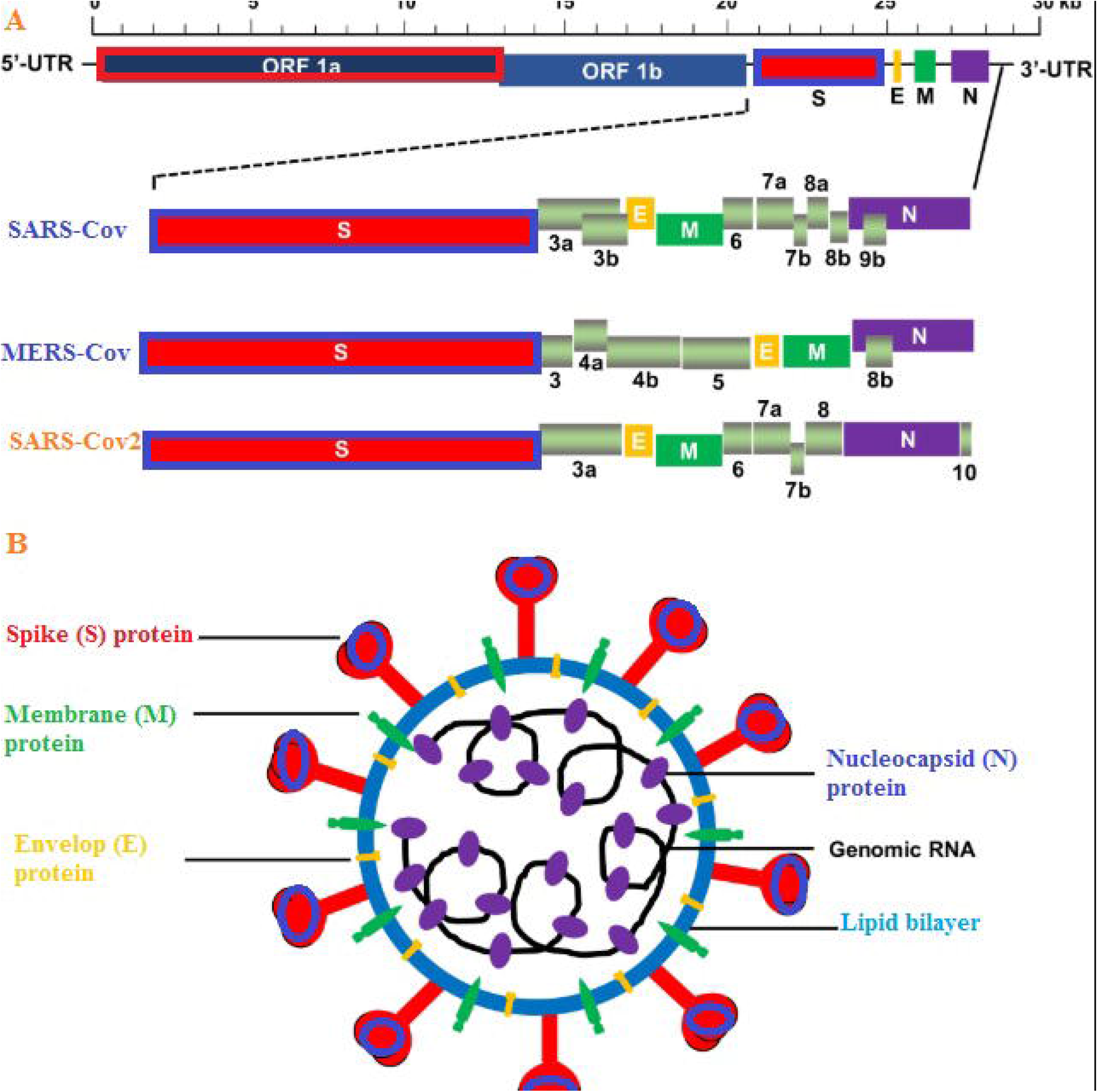
A. Genome organization and protein for SARS Cov2 with respect to SARS Cov, MERS-Cov and SARS-Cov2. B. Arrangements for structural proteins for SARS Cov2

The taxonomy for corona virus includes Subfamily *Othocoronavirinae*, family *Coronaviridae*, order *Nidovirales*, four genera, as *alphacoronavirus (alpha-CoV), betacoronavirus (beta-CoV), gammacoronavirus (gamma-CoV)*, and *deltacoronavirus (delta-CoV)* ^**5**^. Alpha-and beta-CoVs have the ability to infect mammals as bats, pigs, cats, mice, and humans ^**6-11**^. Gamma-and delta-CoVs normally infect birds, while some have the ability to infect mammals^**12-15**^. Considering the importance of the coronavirus spike (S) glycoprotein which facilitates infection through fusion of the viral and cellular membranes with conformational changes^**16**^ and through ACE2 receptors, and consequently, the emerging importance of spike surface protein for the development of drugs and vaccination, we aim to study the phylogeny and genetic variabilities for spike proteins at genomic, amino acid and structural levels.

## Methodology

### Gene sequences from gene bank

Total 562 complete genome sequences of the SARS-CoV-2 strains from the infected individuals are retrieved from the GISAID and NCBI database. We have retrieved some of the nucleotide sequence and amino acid sequence from gene bank, NCBI. For phylogenetic analysis, we retrieved complete genome sequences of SARS Cov2 sequences (multiple sequences from each country) across the globe, pertinent to different continents as Asia, Europe, North America, Africa, South America, Australia. The countries and territories, which are infected by SARS-CoV-2 and share the complete genomes of SARS-COV-2 are Australia (AU), Belgium (BE), Brazil (BR), Canada (CA), Chile (CL), China (CN), Czech Republic (CZ), Denmark (DK), England (UK), Finland (FI), France (FR), Georgia (GE), Germany (DE), Hong Kong (HK), Hungary (HU), India (IN), Ireland (IE), Italy (IT), Japan (JP), Korea (KR), Kuwait (KW), Mexico (MX), Netherlands (NL), New Zealand (NZ), Pakistan (PK), Peru (PR), Russia (RS), Scotland (UK), Singapore (SG), Spain(SP), Switzerland CH), Sweden(SE), Taiwan (TW), Thailand (TH), United Kingdom (UK), Unites States (US), and Vietnam (VN). Gene sequences are shown as in Supplementary file1. We get the spike sequences from the same sequences, retrieved from NCBI gene bank. We retrieved the ACE2 receptor from NCBI protein sequence id <u>NM_021804.3</u> and also the PDB id 1R42.

### Three-dimensional structure prediction and Model quality assessment

PSI-BLAST (http://blast.ncbi.nlm.nih.gov/Blast) was employed for identification of the templates that possessed the highest sequence identity with our target template. PDB structure (3D structure) was developed based on homology modelling of the homologous template structures using PHYRE2 server^**17**^. Molecular visualization tool PyMOL (http://www.pymol.org/) was employed for visualization of 3D structures and its modifications. The Swiss PDB Viewer was used for controlling energy minimization. SAVES (Structural Analysis and Verification Server, http://nihserver.mbi.ucla.edu/SAVES/) was employed for structural evaluation and stereochemical quality assessment of predicted model. The ProSA (Protein Structure Analysis) webserver (https://prosa.services.came.sbg.ac.at/prosa) was employed for refining and validating protein structure^**18**^. It is based on the principle of checking model structural quality with identified errors. The program represents a plot of its residue energies and Z-scores determine the overall quality of the model. TM align software was utilized for the alignment of spike proteins for SARS Cov2 and RMSD estimation to assess the structural differentiation ^**19**^.

### Phylogenetic analysis of SARS Cov 2 world wide

We had studied phylogenetic analysis and detected mutations among the SARS Cov2 sequences spread globally through NJ method using JTT substitution model of MAAFT(https://mafft.cbrc.jp/alignment/server/spool/_phyloio.html) software^**20**^. We attempted to collect complete genome sequences for SARS Cov2 from different countries belonging to different continents.

Considering the wide variability, we had considered phylogenetic tree for different spike proteins for SARS Cov2 studied with MAFFT software.

### Assessment of structural differences between different spike proteins for SARS Cov2

Based on the phylogenetic tree, we identified 20 variants of spike protein existing in the globe. In order to identify the differences, we employed TM align software^**19**^ and assessed RMSD score^**19**^.

### Molecular Docking

Molecular docking is a bioinformatics tool used for *in silico* analysis for the prediction of the binding mode of a ligand with a protein 3D structure. Patch dock is an algorithm for molecular docking based on shape complementarity principle ^**21**^. We subject patchdock analysis of ACE2 receptor with spike protein of SARS Cov2 virus^**21**^. We validated the results with Firedock^**22**^.We had also predicted the binding sites and visualized through Pymol (http://www.pymol.org/).

## Result

### Mutation and sequence variability studies among SARS Cov2 complete genome world wide

High degree of sequence variability has been observed among the SARS Cov2 variants distributed globally. Phylogenetic tree (Fig2) depicts wide variability. However, two distinct clades were observed, but not restricted to infection within a country. An interesting observation was that there exists wide variability among virus strains within a country and even within the same geographical area. We observed multiple strains existing within the same country and also within same geographical location, as in USA (MT88341& MT188340). The other two sequences from USA analyzed even belong to a different clade (MT308704 and MT344956) with genetic variability. Two sequences identified from Kerala state of India and sequenced in the same laboratory depicted enormous genetic variability and they belong to two different clades (MT050493 & MT012098). These conditions are very alarming for effective vaccination or treatment. The detail phylogenetic distribution of SARS COV2 across the globe is being presented in Fig 3, as revealed through GISAID data base (www.gisaid.org) as time progresses.

**Fig 2:**
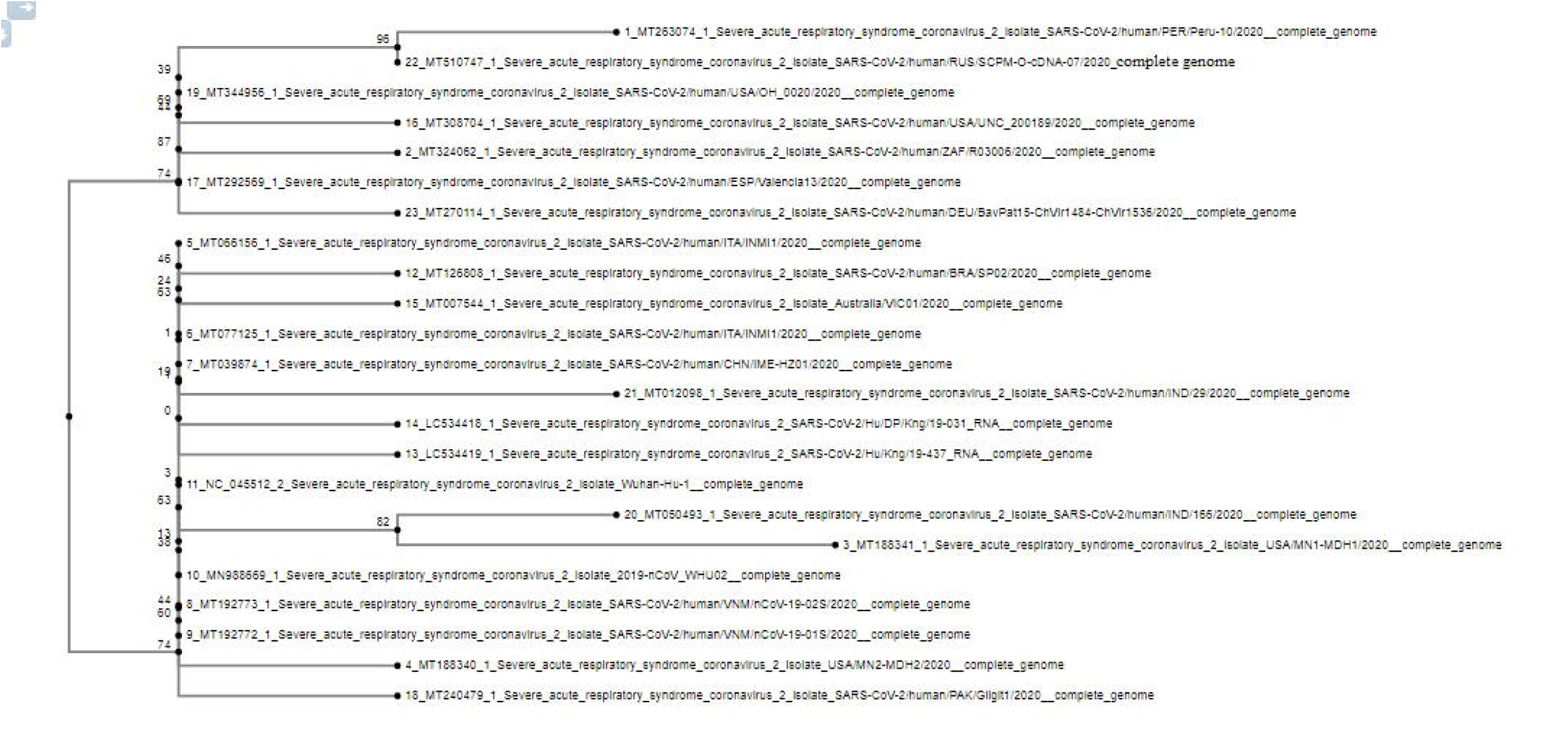
Phylogenetic tree for SARS Cov2-complete genome identified from different countries belonging to all the continents.

**Fig 3:**
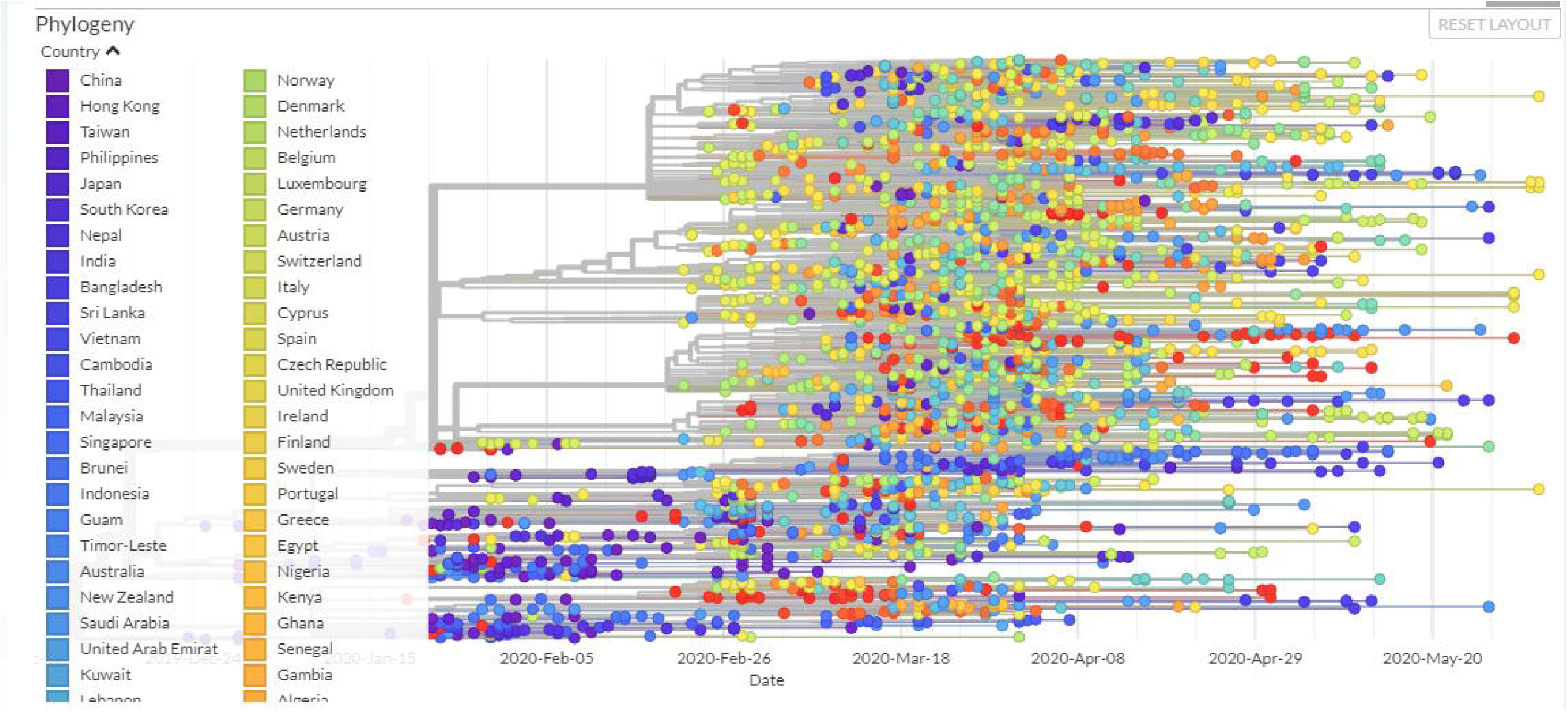
The detail phylogenetic distribution of across the globe as revealed through GISAID data base (www.gisaid.org)

### Mutation and sequence variability studies among SARS Cov2 spike protein world wide

Molecular phylogeny for different variants for spike proteins were depicted through phylogenetic tree in Fig 4. Out of 20 different spike proteins identified, 7 types were clustered together, indicating genetic closeness. They may be regarded as the dominant type. Phylogenetic tree was constructed which reveal spike protein 1 (QIS60288.1), 2 (QIZ15537), 13 (QIV64989), 14(QIU78707), 16(QJA17668), 19(QJZ27839) and 20 (QJW69307) were clustered together, indicating genetic similarity and may be regarded as dominant type.

**Fig 4:**
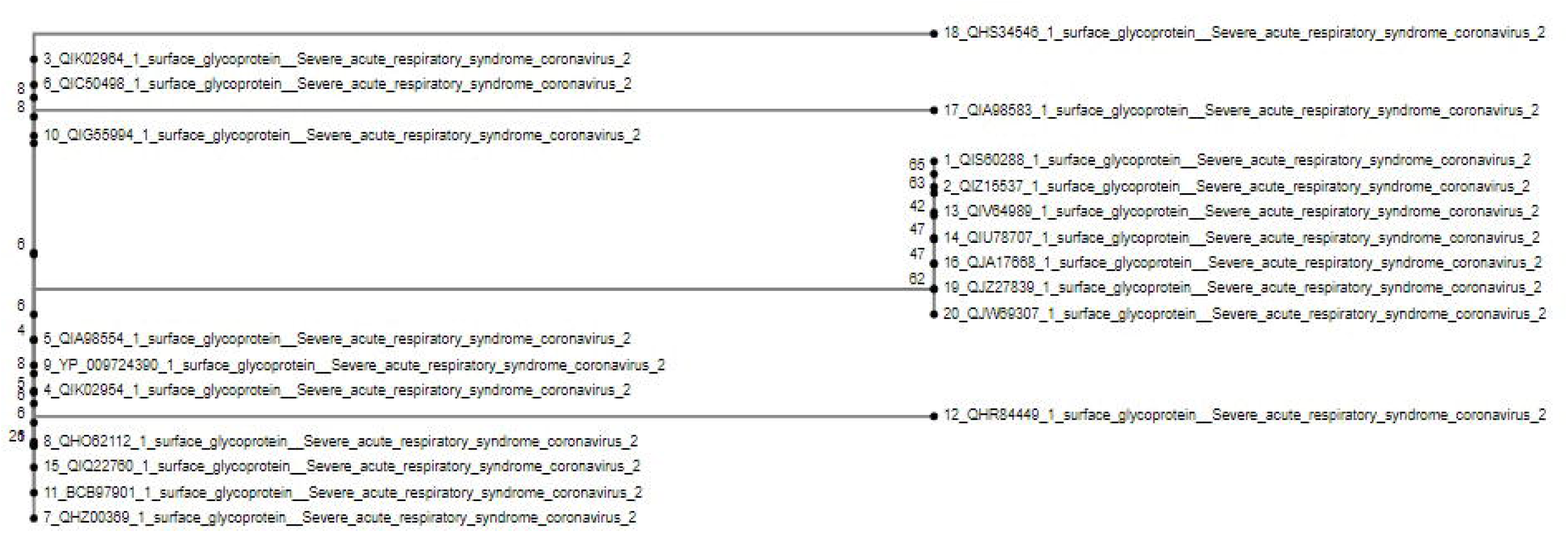
Phylogenetic tree for spike variants of SARS Cov2 distributed globally

Alignment of different variants of 3D structure of spike protein for SARS Cov2 were compared and RMSD listed in Table 1. We have visualized alignment report of genetically distant variants with Spike protein1, considering it as dominant type. Certain important novel non-synonymous mutations have been identified among the variants of spike protein leading to amino acid changes as S247R, R408I, G612D, A930V and deletion detected at amino acid position 144 (Fig 5). R408I lies within Peptidase S domain. 3D structural alignments are revealed with respect to dominant type spike 1 (Fig 6)

**Table 1:** RMSD (Random mean square deviation) value due to alignment of 3D structure for different variants of Spike protein with the dominant type (Spike 1) by TM align

|  |  |  |  |  |  |
| --- | --- | --- | --- | --- | --- |
|  | 1 |  | 1 |  | 1 |
| 3 | 3.51 | 8 | 3.19 | 15 | 4.44 |
| 4 | 4.45 | 9 | 4.37 | 17 | 2.25 |
| 5 | 4.14 | 10 | 4.14 | 18 | 7.28 |
| 6 | 3.28 | 11 | 3.28 | 20 | 3.09 |
| 7 | 3.46 | 12 | 2.45 |  |  |

**Fig 5:**
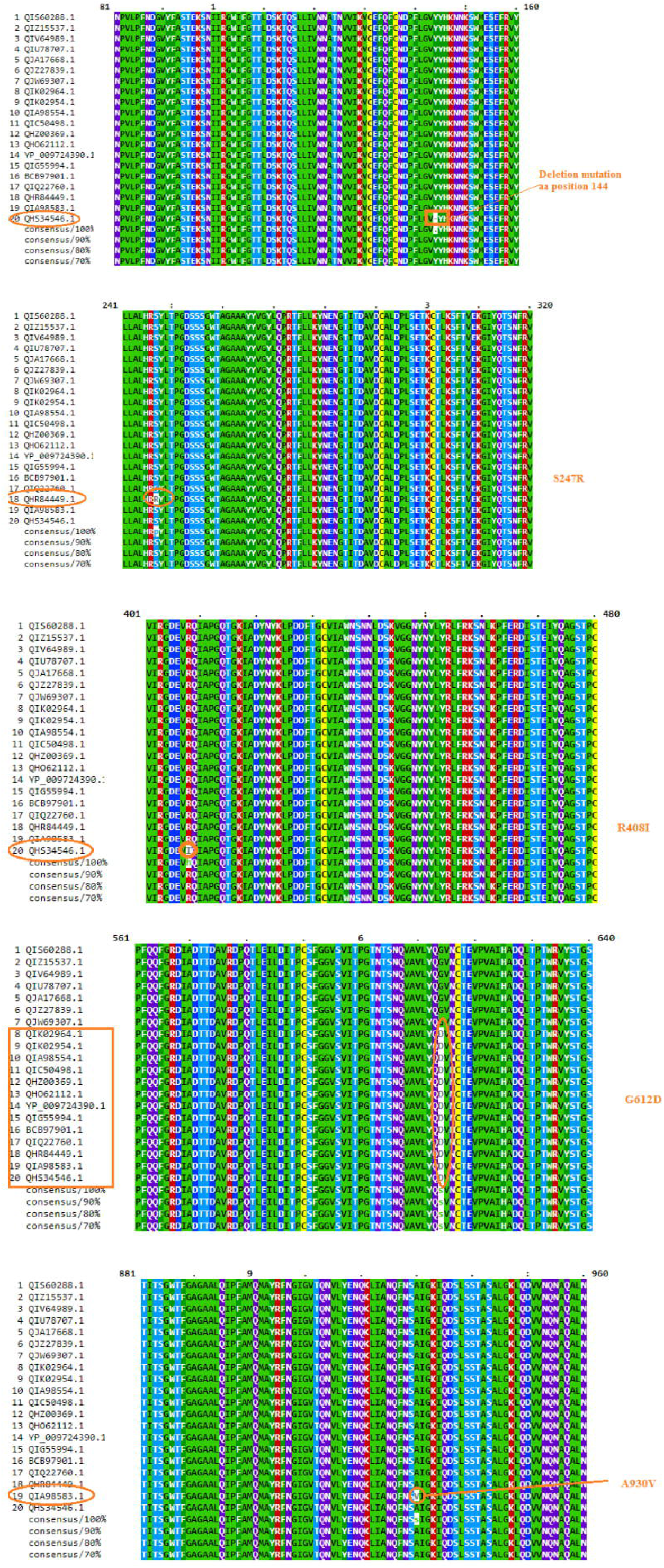
Alignment for different variants of Spike protein with the dominant type (Spike 1). Amino acid variations identified.

**Fig 6:**
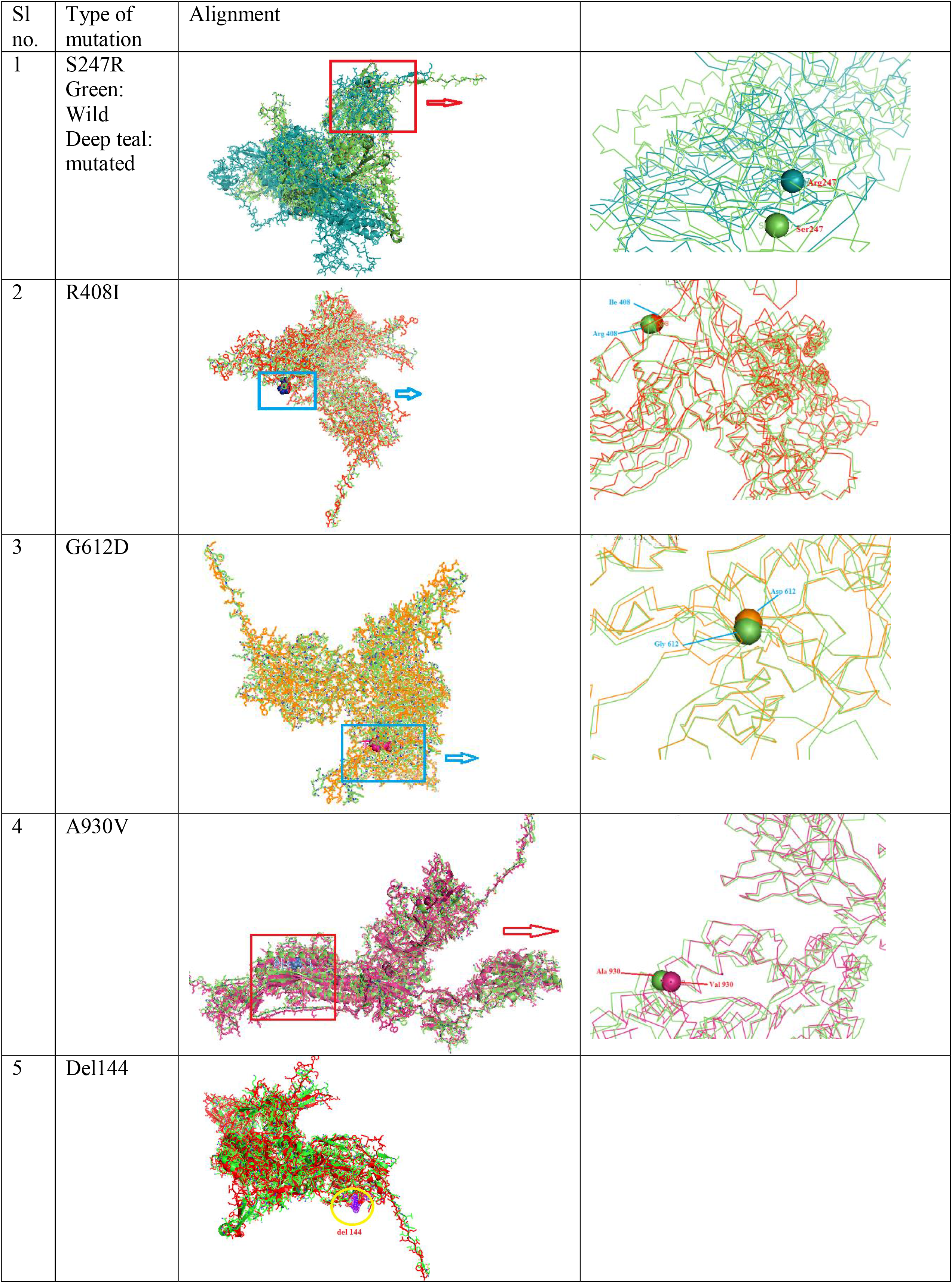
Alignment of 3D structure of dominant type spike 1 with mutated variants of spike protein of SARS Cov2

However, the alignment studies reveal structural differences as revealed through RMSD. The structural differences were clearly observed with maximum variability observed in variant 1 vs. 4 (RMSD:4.45) and 1vs. 15 (RMSD: 4.44). We had considered variant1 as representative for the dominant variant. Maximum variability observed for variant 4 (from US) and variant 5 from Pakistan (patient had travel history to Iran, and returned to Pakistan 5 days before sample was collected).

### Molecular Docking of ACE2 receptor with spike protein

We assessed the binding ability of ACE2 receptor with dominant spike protein1 with binding sites detected. (Fig 7a &7b.).We identified molecular docking of ACE2 receptor (blue) with spike protein(red) for SARS Cov2 virus. We used spike protein1 for binding with ACE2 receptor as in Fig 7a, 7b. The binding domains for ACE2 with spike protein are being identified as Ile54, Gly337, Val 293, Gln305, Lys363, Asn394, Lys309, Asn149 as represented as yellow sphere(Fig 7a).

**Fig 7:**
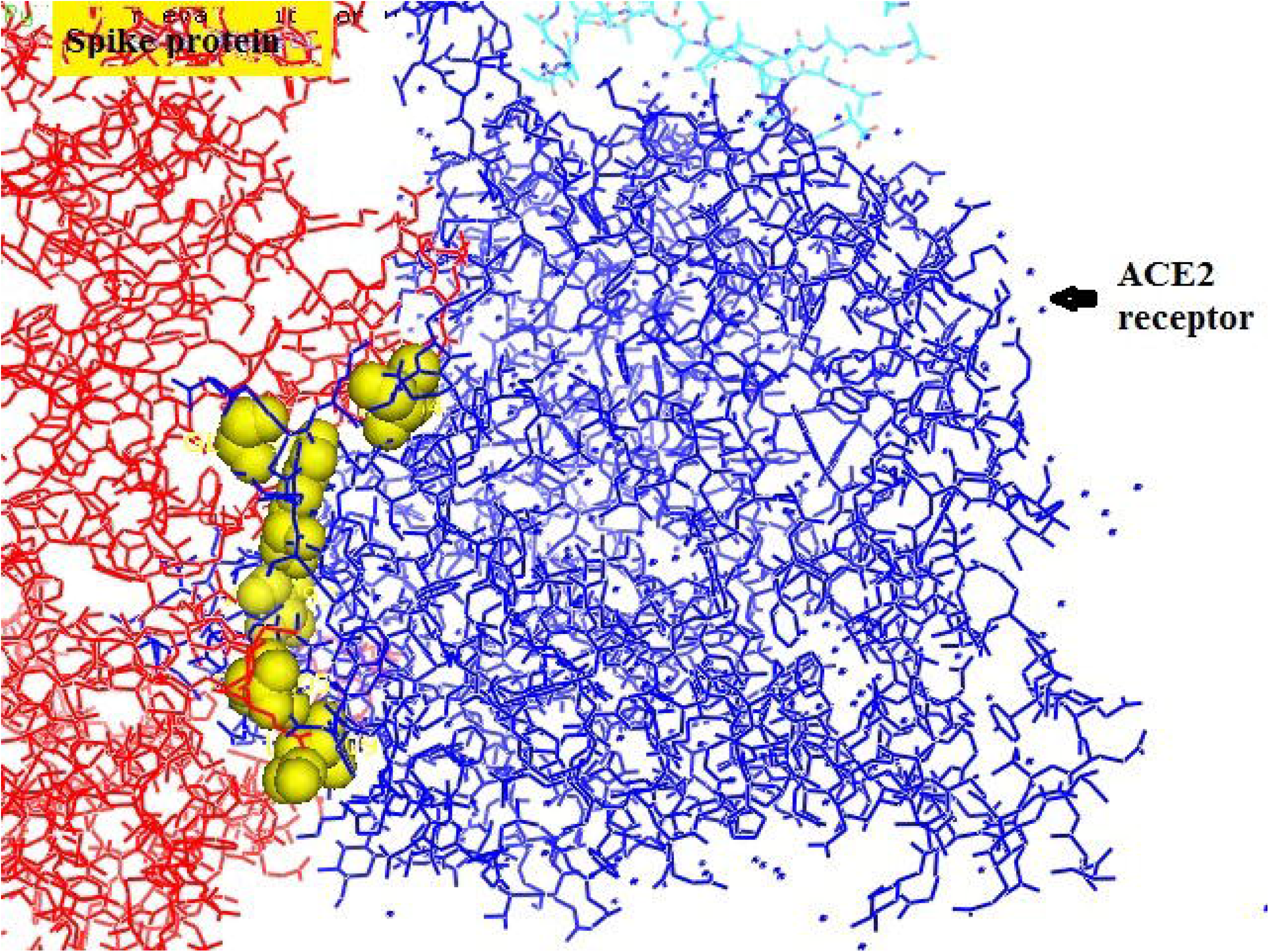
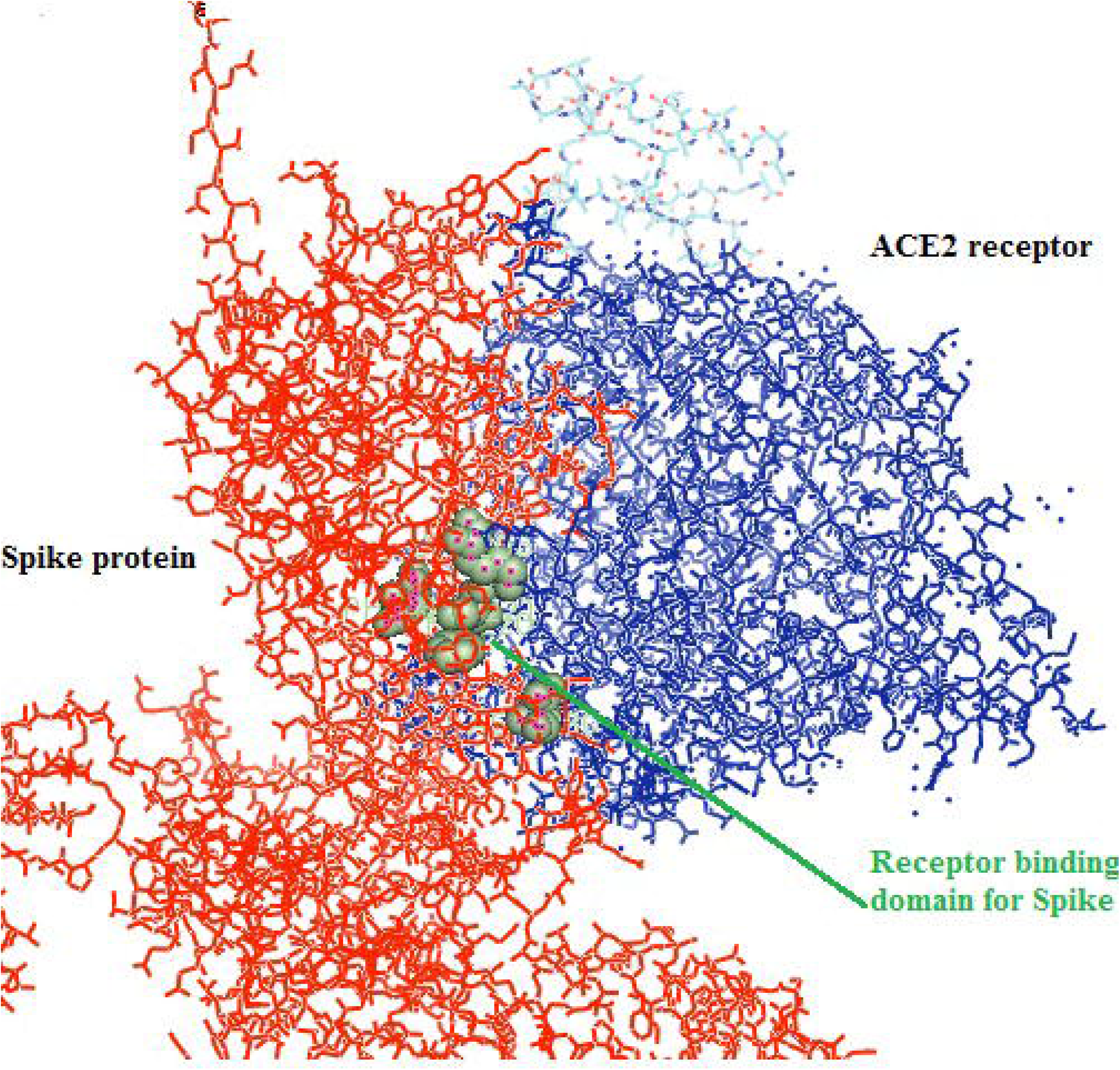
Molecular docking of spike with ACE2 receptor, a. Prediction of binding site of ACE2 with Spike protein b. Prediction of Binding domain of Spike protein withACE2 receptor.

Simultaneously, we also predicted the receptor binding domains for spike protein1 with ACE2 receptor as Ser221, Phe220, Phe32, Asn 280, Lys206 (Fig 7b).

## Discussion

We identified a wide genetic variabilities among the SARS Cov2 complete genome sequences distributed worldwide across different countries belonging to different continents. Although, primarily two clades were identified, but they are not restricted to a particular geographical area. Even within a country, within the same geographical area, genetic differences were observed. In our study, we observed sufficient genetic variabilities among the SARS Cov2 virus within two different samples collected from Kerala, India. In the same way, different strains were identified from the same state of USA. Even within the same country USA, different strains belong to two different clades. Evolutionary studies for SARS Cov2 have been conducted by a number of workers, but due to rapidly evolving virus, molecular phylogeny varies frequently. Initial report indicates two major types^**23**^, three types ^**24**^ followed by 11 subtypes^**25**^. It is suggested that O type as the ancestral type first evolved from China, Wuhan outbreak, while A2 type as the dominant one, characterized by a non-synonymous D614G mutation at the junction of S1-RBD and S2 of the SARS-CoV-2 spike (S) protein^**25**^. This high genetic variability restricts the development of satisfactory vaccines. Lack of genetic uniformity within a particular region, makes the vaccination process more complicated. The possibilities of transmission of SARS Cov2 through animals (bat, pangolin, reptiles, ferret, pet animals as dog, cat, clinical trial in monkey) with zoonotic importance have been well discussed^**26**^.

Considering the importance of surface glycoprotein or spike protein in attachment to ACE receptor, during the entry in host, we identified 20 different spike variants numbered as (1-20). Phylogenetic tree revealed seven spike proteins (1,2,13,14, 16 19, 20) to be genetically similar and clustered together. However there still exists certain differences in 3D structures among these spike proteins. It is interesting to note that spike variant18 has a different structure compared to others. Such differences have also been detected by researchers. Reports are available that the coronavirus spike (S) glycoprotein fuses with host cellular membranes via ACE receptor by conformation changes of the spike^**16**^. This has been illustrated through biochemical characterization, limited proteolysis, mass spectrometry, and single-particle EM ^**16**^. It has been reported that that the stability of SARS-CoV-2 Spike protein is less than SARS-CoV S ^27^. We detected certain novel mutations at S247R, R408I, G612D, A930V and del 144, one involved with important domain of peptidase S. We presume that the variabilities in spike protein have arisen due to non-synonymous mutations in spike protein distributed world wide. Such wide variabilities in ssRNA virus is very common as we detected in avian influenza virus^**28**^. It has been reported that SARS-CoV-2 binds with human angiotensin converting enzyme 2 (ACE2) ^1^ as a receptor and enter the human lung. The spike (S) protein acts through receptor binding and membrane fusion ^29^. The spike protein of coronaviruses has two functional domains – S1, involved with receptor binding, and S2 domain, causes cell membrane fusion^30^.

We identified the receptor binding domain sites for ACE2 receptor as Ser221, Phe220, Phe32, Asn 280, Lys206. However others have identified five key residues in the receptor binding domain enable efficient binding of SARS-CoV-2 to human ACE2; these are Asn439, Asn501, Gln493, Gly485 and Phe486^**38**^. The variation might occur due to wide variability in spike proteins identified. Another mutation, A23403G, located in the gene encoding the spike glycoprotein results in an amino acid change (D614G) from aspartic acid to glycine. Although the effect of the D614G mutation is not clear, the location of this mutation was reported in the S1-S2 junction near the furin recognition site (R667) for the cleavage of S protein. This site was observed to be essential for the entry of the virion into the host cell^30^. In this current study, we had also identified four non-synonymous mutations and one site for deletion. It is important to note that R408I lies within Peptidase S domain, which might be responsible for variation in 3D structures of the spikes leading to greater RMSD. This might aid in altered pathogenicity affecting recognition and binding of host receptor. With the help of SARS-CoV-2 S protein pseudovirus system, it was proved that human angiotensin converting enzyme 2 (hACE2) is the receptor for SARS-CoV-2. It has been reported that SARS-CoV-2 gets its entry through 293/hACE2 cells mainly through endocytosis. Factors such as PIKfyve, TPC2, and cathepsin L are reported to be critical for the entry^**27**^. The wide variabilities in spike protein leads to different domain binding site for ACE2 receptor. This may also be one of the reason for difference in susceptibility to infection. Other researchers identified polymorphism of ACE2 receptor and regard it to be responsible for varying susceptibility^31^.

An important subunit observed with greater affinity and binding ability was S (surface glycoprotein molecule) for most of the immune response molecule studied. Considering its importance studied earlier, it has been considered as an effective antigen molecule for subunit vaccine production^**32**^. Spike protein of human CoVs help in viral entry into host cells leading to membrane fusion, causing viral infection^**32,33**^. The S protein is a class I viral protein, that cleave into two functional subunits, designated as S1 and S2. S1 is an amino-terminal subunit and S2 is a carboxyl-terminal subunit. The function of S1 subunit is virus–host cell receptor binding, whereas the S2 subunit is responsible for virus–host membrane fusion^**32**^.

Two major domains are present in S1, termed as an N-terminal domain (NTD) and a C-terminal domain (CTD). NTDs aid in sugar binding^**32**^, whereas CTDs act as protein receptor. S1 subunit is responsible for binding host receptors or function as receptor-binding domains (RBDs) ^**32**^. The viral membrane protein interacts with host cellular membrane protein and ultimately enters the host. The infection is initiated as S1 subunit recognize receptor or sugar moiety on the cell surface. As the next step, the S protein undergoes conformational changes, leading to membrane fusion by the S2 region^**32**^. Finally, the viral genetic particles are inserted or delivered into the host cell by the fusion core^**32,34**^. Reports indicate the existence of spike glycoprotein as metastable prefusion trimer as characterized by its structure^**32**^. The structural and functional basis of receptor binding site for SARS Cov2 have been studied^**35,36**^. We observed spike protein 18 as completely different from others, which may occur due to the conformational changes of spike.

The receptor binding domain (CTD) of the trimeric spike protein exists in two states -standing or lying. ACE2 receptor are able to bind to standing RBD, particularly in the receptor-binding motif (RBM), aiding the RBD to exist in the “standing” state^**32**^. Along with human ACE2, SARS-CoV2 S protein have the ability to bind to palm civet and mouse ACE2s^**35,36**^. It has been reported that mutations in the RBD of S1 subunit are essential for cross-species transmission of SARS-CoV^**32**^. Like SARS-CoV, 2019-nCoV also employs ACE2 as its cellular receptor for its entry in host cells^**32**^.

We observed ACE2 receptor was able to bind effectively with different identified variants with non-synonymous mutations *in silico*, thus initiates infection. This might be due to the fact that identified the samples were from virulent strains of SARS Cov2. However, polymorphism of ACE2 receptor had also been identified^31^, that might also explain the variabilities in pathogenicity in SARS Cov2.

### Conclusion

We identified wide variabilities in SARSCov2 virus distributed world wide and even within same geographical region as revealed through phylogenetic tree. Out of 20 different variants of spike proteins identified, seven were observed to be genetically similar and may be regarded as dominant type, may be helpful for vaccination or drug development. Five novel mutations detected in spike proteins as S247R, R408I, G612D, A930V and del 144,when R408I involved with important domain of peptidase S. ACE2 receptor bind with the variants of spike proteins.

## Key points

☛Wide variability in the spike protein variants (binds to ACE2 receptor) of SARS Cov2 identified globally.
☛ Dominant type of spike protein identified
☛ Novel non-synonymous and deletion mutations identified for spike protein w.r.t. dominant variant spike1
☛ RMSD values ranges from 4.45 to 2.25 for the dominant variant spike1 w.r.t other spike protein 3D structure.
☛ Scope for effective vaccine and drug development with the dominant type of spike protein.

## Notes

### Competing Interest Statement

The authors have declared no competing interest.

## References

1. Lu R, Zhao X, Li J, Niu P, Yang B, Wu H et al. Genomic characterisation and epidemiology of 2019 novel coronavirus: implications for virus origins and receptor binding. Lancet 2020; 395: 565–74.

2. Bhattacharya, C., Das, C., et al. Global spread of SARS Cov2 subtype with Spike Protein Mutation D614G is shaped by Human genomic variations that regulate expression of TMPRSS2 and MX1 genes. bioRxiv preprint doi: https://doi.org/10.101/2020.05.04.075911.

3. Verity R, Okell LC, Dorigatti I, Winskill P, Whittaker C, Imai N, et al. Estimates of the severity of coronavirus disease 2019: a model-based analysis. Lancet Infect Dis. Published Online March 30, 2020. https://doi.org/10.1016/S1473-3099(20)30243-7.

4. World-Health-Organization Update 49 - SARS case fatality ratio, incubation period. Available online: https://www.who.int/csr/sars/archive/2003_05_07a/en/ (accessed on 9 April 2020).

5. King, A. M. Q., Lefkowitz, E. J., Mushegian, A. R., Adams, M. J., Dutilh, B. E., Gorbalenya, A. E., et al.. Changes to taxonomy and the international code of virus classification and nomenclature ratified by the international committee on taxonomy of viruses (2018). Arch. Virol. 163, 2601–2631. doi: 10.1007/s00705-018-3847-1

6. Kusanagi, K., Kuwahara, H., Katoh, T., Nunoya, T., Ishikawa, Y., Samejima, T., et al. (1992). Isolation and serial propagation of porcine epidemic diarrhhea virus in cell-cultures and partial characterization of the isolate. J. Vet. Med. Sci. 54, 313–318. doi: 10.1292/jvms.54.313

7. Li, W., Shi, Z., Yu, M., Ren, W., Smith, C., Epstein, J. H., et al. (2005b). Bats are natural reservoirs of SARS-like coronaviruses. Science 310, 676–679. doi: 10.1126/science.1118391

8. Poon, L. L., Chu, D. K., Chan, K. H., Wong, O. K., Ellis, T. M., Leung, Y. H., et al. (2005). Identification of a novel coronavirus in bats. J. Virol. 79, 2001–2009. doi: 10.1128/jvi.79.4.2001-2009.2005

9. Drexler, J. F., Corman, V. M., and Drosten, C. (2014). Ecology, evolution and classification of bat coronaviruses in the aftermath of SARS. Antiviral Res. 101, 45–56. doi: 10.1016/j.antiviral.2013.10.013

10. Pedersen, N. C. (2014). An update on feline infectious peritonitis: virology and immunopathogenesis. Vet. J. 201, 123–132. doi: 10.1016/j.tvjl.2014.04.017

11. Cui, J., Li, F., and Shi, Z. L. (2019). Origin and evolution of pathogenic coronaviruses. Nat. Rev. Microbiol. 17, 181–192. doi: 10.1038/s41579-018-0118-9

12. Woo, P. C., Lau, S. K., Lam, C. S., Lai, K. K., Huang, Y., Lee, P., et al. (2009a). Comparative analysis of complete genome sequences of three avian coronaviruses reveals a novel group 3c coronavirus. J. Virol. 83, 908–917. doi: 10.1128/jvi.01977-08

13. Woo, P. C., Lau, S. K., Lam, C. S., Lau, C. C., Tsang, A. K., Lau, J. H., et al. (2012). Discovery of seven novel mammalian and avian coronaviruses in the genus deltacoronavirus supports bat coronaviruses as the gene source of alphacoronavirus and betacoronavirus and avian coronaviruses as the gene source of gammacoronavirus and deltacoronavirus. J. Virol. 86, 3995–4008. doi: 10.1128/jvi.06540-11

14. Woo, P. C., Lau, S. K., Lam, C. S., Tsang, A. K., Hui, S. W., Fan, R. Y., et al. (2014). Discovery of a novel bottlenose dolphin coronavirus reveals a distinct species of marine mammal coronavirus in gammacoronavirus. J. Virol. 88, 1318–1331. doi: 10.1128/jvi.02351-13

15. Ma, Y., Zhang, Y., Liang, X., Lou, F., Oglesbee, M., Krakowka, S., et al. (2015). Origin, evolution, and virulence of porcine deltacoronaviruses in the United States. mBio 6:e00064. doi: 10.1128/mBio.00064-15

16. Walls, A.C., Tortorici, M.A., Snijdera, J., Xionga, X., Boschd, B.J., Rey, F.A. and Veeslera, D. Tectonic conformational changes of a coronavirus spike glycoprotein promote membrane fusion. PNAS | October 17, 2017 | vol. 114 | no. 42 | 11157–11162. /doi/10.1073/pnas.1708727114

17. Kelley L. et al. The Phyre2 web portal for protein modeling, prediction and analysis. Nature Protocols; 10, 845-854.doi.org/10.1038/nprot.2015.053 (2015).

18. Wiederstein M., et al. ProSA-web: interactive web service for the recognition of errors in three-dimensional structures of proteins, Nucleic Acids Research, Volume 35, Issue suppl_2, Pages W407–W410, https://doi.org/10.1093/nar/gkm290 (2007).

19. Zhang Y, et al. TM-align: A protein structure alignment algorithm based on TM-score, Nucleic Acids Research, 33: 2302–2309 (2005).

20. Katoh K., et al. MAFFT multiple sequence alignment software version 7: improvements in performance and usability. Molecular biology and evolution, 30(4), 772–780. https://doi.org/10.1093/molbev/mst010 (2013).

21. Duhovny, D.S., et al. PatchDock and SymmDock: servers for rigid and symmetric docking. Nucleic Acids Res. 2005 1; 33(Web Server issue): W363–W367. doi: 10.1093/nar/gki481 (2005).

22. Mashiach E., et al. FireDock: a web server for fast interaction refinement in molecular docking. Nucleic acids research, 36(Web Server issue), W229–W232. https://doi.org/10.1093/nar/gkn186 (2008).

23. Tang, X., Wu, C., Li, X., Song, Y., Yao, X., Wu, X. et al., On the origin and continuing evolution of SARS-CoV-2. National Science Review, nwaa036, 2020. https://doi.org/10.1093/nsr/nwaa036)(in press)

24. Forster, P., Forster, L., Renfrew, C., Forster, M. Phylogenetic networkanalysis of SARS-CoV-2 genomes. Proc. Natl.Acad. Sci.USA 2020 (In press)

25. Biswas, N. and Majumder, P. 2020. Analysis of RNA sequences of 3636 SARS Cov2 collected from 55 countries reveals selective sweep of one virus type. Indian Journal of Medical research. In press.

26. Mahdy, M.A.A. 2020. An overview of SARS Cov2 and Animal infection. Preprints. Doi.10.20944/preprints202004

27. Ou, X., Liu, Y., Lei, X., Li, P., Mi, D., Ren, L., Guo, L., Guo, R., Chen, T., Hu, J., Xiang, Z., Mu, Z., Chen, X., Chen, J., Hu, K., Jin, Q., Wang, J., & Qian, Z. Characterization of spike glycoprotein of SARS-CoV-2 on virus entry and its immune cross-reactivity with SARS-CoV. Nature communications. https://doi.org/10.1038/s41467-020-15562-9

28. Pal, A., Pal, Ab and Baviskar, P. 2020. RIGI, TLR7 and TLR3 confer better antiviral resistance to indigenous ducks against Avian influenza –scope to develop a novel therapeutic approach to prevent another pandemic Flu. Antiviral research (submitted).

29. Li F. Structure, function, and evolution of coronavirus spike proteins. Annu Rev Virol 2016; 3: 237–61.

30. Follis KE, York J, Nunberg JH. Furin cleavage of the SARS coronavirus spike glycoprotein enhances cell– cell fusion but does not affect virion entry. Virology 2006; 350: 358–69.

31. Devaux, C.A., Rolain, J.M., Raoult, D. ACE2 receptor polymorphism: Susceptibility to SARS-CoV-2, hypertension, multi-organ failure, and COVID-19 disease outcome. Journal of Microbiology, Immunology and Infection. Volume 53, Issue 3, June 2020, Pages 425–435.

32. Wang, N., Shang, J., Jiang, S., Du, L. Subunit vaccine against emerging pathogenic Human coronavirus. Frontiers Microbiology. 2020. https://doi.org/10.3389/fmicb.2020.00298.

33. Du, L., He, Y., Zhou, Y., Liu, S., Zheng, B. J., and Jiang, S. (2009). The spike protein of SARS-CoV - a target for vaccine and therapeutic development. Nat. Rev. Microbiol. 7, 226–236. doi: 10.1038/nrmicro2090

34. He Y, Zhou Y, Liu S, et al. Receptor-binding domain of SARS-CoV spike protein induces highly potent neutralizing antibodies: implication for developing subunit vaccine. Biochem Biophys Res Commun 2004; 324: 773–81.

35. Shang, J., Ye, G., Shi, K. et al. Structural basis of receptor recognition by SARS-CoV-2. Nature 581, 221–224 (2020). https://doi.org/10.1038/s41586-020-2179-y

36. Wang et al., 2020, Cell 181, 894–904 May 14, 2020 ^a^ 2020 Elsevier Inc. https://doi.org/10.1016/j.cell.2020.03.045

