## Supplementary file 1 for "Wide variabilities identified among spike proteins of SARS Cov2 globally-dominant variant identified"

GISAID ID

FR|HF1805|EPI_ISL_414628|2020-03-02

FR|IDF2075|EPI_ISL_415650|2020-03-02

FR|IDF2075|EPI_ISL_415693|2020-03-02

FR|B2348|EPI_ISL_416511|2020-03-07

FR|HF2174|EPI_ISL_415654|2020-03-09

DE|NRW-01|EPI_ISL_413488|2020-02-28

UK|SHEF-BFCB1|EPI_ISL_416730|2020-03-03

UK|SHEF-BFCDF|EPI_ISL_416732|2020-03-03

UK|SHEF-BFCFD|EPI_ISL_416734|2020-03-09

UK|SHEF-BFCC0|EPI_ISL_416731|2020-03-03

UK|EDB003|EPI_ISL_415640|2020-03-08

UK|EDB004|EPI_ISL_415629|2020-03-10

UK|CVR05|EPI_ISL_414027|2020-02-26

UK|CVR07|EPI_ISL_415630|2020-03-10

UK|SHEF-BFCEE|EPI_ISL_416733|2020-03-07

UK|CVR10|EPI_ISL_415631|2020-03-10

UK|SHEF-BFD45|EPI_ISL_416739|2020-03-09

UK|SHEF-BFD54|EPI_ISL_416740|2020-03-03

DE|NRW-06|EPI_ISL_414505|2020-03-04

DE|NRW-09|EPI_ISL_414509|2020-02-28

UK|200990724|EPI_ISL_414006|2020-02-28

UK|200690756|EPI_ISL_414044|2020-02-08

UK|200641094|EPI_ISL_414040|2020-02-05

UK|200690245|EPI_ISL_414041|2020-02-08

UK|200690300|EPI_ISL_414042|2020-02-08

UK|200690306|EPI_ISL_414043|2020-02-07

IT|SSI-101|EPI_ISL_415646|2020-03-03

FR|HF2234|EPI_ISL_416495|2020-03-10

FR|IDF2278|EPI_ISL_416499|2020-03-11

CH|1000477757|EPI_ISL_413021|2020-02-29

CH|1000477806|EPI_ISL_413024|2020-02-29

CH|GE3895|EPI_ISL_413997|2020-02-26

CH|GE1422|EPI_ISL_415454|2020-02-29

CH|GE1402|EPI_ISL_415700|2020-02-28

CH|AG7120|EPI_ISL_415457|2020-02-29

CH|SZ1417|EPI_ISL_415702|2020-03-02

CH|BE6651|EPI_ISL_415456|2020-02-29

CH|GE9586|EPI_ISL_414022|2020-02-27

CH|BL0902|EPI_ISL_414021|2020-02-27

CH|BS0914|EPI_ISL_415701|2020-03-02

CH|GE0199(2020|EPI_ISL_415455|2020-02-28

CH|TI2045|EPI_ISL_415703|2020-03-01

CH|GE8102|EPI_ISL_415458|2020-03-01

CH|GR2988|EPI_ISL_415698|2020-02-27

CH|GR3043|EPI_ISL_415699|2020-02-27

CH|VD0503|EPI_ISL_415459|2020-02-29

CH|VD5615|EPI_ISL_414023|2020-03-01

FR|GE1583|EPI_ISL_414623|2020-02-25

FR|GE1583|EPI_ISL_414600|2020-02-26

DE|BavPat2|EPI_ISL_414520|2020-03-02

SE|01|EPI_ISL_411951|2020-02-07

FR|B2340|EPI_ISL_416507|2020-03-05

FR|IDF0515-isl|EPI_ISL_410984|2020-01-29

FR|IDF0515|EPI_ISL_408430|2020-01-29

FR|IDF0571|EPI_ISL_411218|2020-02-02

DE|BavPat3|EPI_ISL_414521|2020-03-02

FR|IDF0372|EPI_ISL_406596|2020-01-23

FR|IDF0386-islP3|EPI_ISL_411220|2020-01-28

FR|IDF0386-islP1|EPI_ISL_411219|2020-01-28

FR|IDF0372-isl|EPI_ISL_410720|2020-01-23

IT|SPL1|EPI_ISL_412974|2020-01-29

FR|IDF0626|EPI_ISL_408431|2020-01-29

FR|HF1993|EPI_ISL_414637|2020-03-04

FR|HF1870|EPI_ISL_414629|2020-03-03

FR|B2344|EPI_ISL_416509|2020-03-06

FR|GE1977|EPI_ISL_414632|2020-03-04

FR|IDF1980|EPI_ISL_414633|2020-03-04

FR|BFC2147|EPI_ISL_415652|2020-03-05

FR|BFC2147|EPI_ISL_415695|2020-03-05

FR|HF1988|EPI_ISL_414635|2020-03-04

FR|HF1995|EPI_ISL_414638|2020-03-04

FR|N1620|EPI_ISL_414601|2020-02-27

IT|CDG1|EPI_ISL_412973|2020-02-20

IT|UniSR1|EPI_ISL_413489|2020-03-03

FR|HF1684|EPI_ISL_414626|2020-02-29

CH|AG0361|EPI_ISL_413999|2020-02-27

CH|GE4984|EPI_ISL_415708|2020-03-07

CH|GE3121|EPI_ISL_414019|2020-02-27

CH|GE5373|EPI_ISL_414020|2020-02-27

CH|TI9486|EPI_ISL_413996|2020-02-24

FR|HF2196|EPI_ISL_416493|2020-03-08

CH|BE2536|EPI_ISL_415704|2020-03-04

CH|GE4135|EPI_ISL_415705|2020-03-06

CH|GE062072020|EPI_ISL_415706|2020-03-06

CH|GE6679|EPI_ISL_415707|2020-03-08

FR|B2330|EPI_ISL_416502|2020-02-26

UK|200990723|EPI_ISL_414012|2020-02-27

CH|1000477797|EPI_ISL_413023|2020-02-29

DE|Baden-Wuerttemberg-1|EPI_ISL_412912|2020-02-25

CH|1000477796|EPI_ISL_413022|2020-02-29

UK|200990002|EPI_ISL_414522|2020-02-28

UK|200960041|EPI_ISL_414008|2020-02-27

FR|B2349|EPI_ISL_416512|2020-03-07

UK|PHW32|EPI_ISL_415920|2020-03-12

UK|PHW37|EPI_ISL_415657|2020-03-12

UK|Sheff01|EPI_ISL_414500|2020-03-03

FR|HF1871|EPI_ISL_414630|2020-03-03

UK|PHW27|EPI_ISL_415655|2020-03-05

FR|HF1795|EPI_ISL_414627|2020-03-02

FR|HF2039|EPI_ISL_415649|2020-03-05

FR|HF2039|EPI_ISL_415692|2020-03-05

IT|SSI-104|EPI_ISL_415648|2020-03-03

FR|B2351|EPI_ISL_416513|2020-03-07

FR|IDF2256|EPI_ISL_416498|2020-03-11

FR|HF2237|EPI_ISL_416496|2020-03-10

FR|PL1643|EPI_ISL_414625|2020-02-26

UK|200991076|EPI_ISL_414524|2020-03-01

FR|HF2239|EPI_ISL_416497|2020-03-10

FR|N1620|EPI_ISL_414624|2020-02-26

FR|BFC2094|EPI_ISL_415651|2020-03-05

FR|BFC2094|EPI_ISL_415694|2020-03-05

FR|B2346|EPI_ISL_416510|2020-03-06

FR|B2337|EPI_ISL_416506|2020-03-03

FR|B2336|EPI_ISL_416505|2020-03-02

DE|BavPat1|EPI_ISL_406862|2020-01-28

UK|201040141|EPI_ISL_414526|2020-03-03

UK|201040081|EPI_ISL_414525|2020-03-02

UK|200990660|EPI_ISL_414523|2020-02-27

FR|N2223|EPI_ISL_416494|2020-03-04

FR|GE1973|EPI_ISL_414631|2020-03-04

UK|PHW1|EPI_ISL_413555|2020-02-27

UK|02|EPI_ISL_407073|2020-01-29

UK|01|EPI_ISL_407071|2020-01-29

UK|200940527|EPI_ISL_414005|2020-02-25

UK|09c|EPI_ISL_412116|2020-02-09

FR|B2334|EPI_ISL_416503|2020-03-01

UK|200990006|EPI_ISL_414011|2020-02-26

FR|IDF0373|EPI_ISL_406597|2020-01-23

UK|200990725|EPI_ISL_414007|2020-02-28

UK|200960515|EPI_ISL_414009|2020-02-25

NL|ZuidHolland_16|EPI_ISL_414558|2020-03-04

NL|NA21|EPI_ISL_415478|2020-03-10

NL|NA35|EPI_ISL_415492|2020-03-10

NL|NoordBrabant56|EPI_ISL_415512|2020-03-09

GE|Tb-54|EPI_ISL_415641|2020-02-27

FI|FIN0303B|EPI_ISL_413603|2020-03-03

NL|Utrecht18|EPI_ISL_415527|2020-03-09

NL|NA29|EPI_ISL_415486|2020-03-10

NL|ZuidHolland_8|EPI_ISL_414468|2020-03-06

NL|NA34|EPI_ISL_415491|2020-03-10

NL|NoordBrabant_6|EPI_ISL_414451|2020-03-06

NL|Rotterdam_1363790|EPI_ISL_413582|2020-03-01

NL|Hardinxveld_Giessendam_1364806|EPI_ISL_413573|2020-03-02

NL|Naarden_1364774|EPI_ISL_413577|2020-03-02

NL|Rotterdam_1364040|EPI_ISL_413583|2020-03-02

GE|Tb-537|EPI_ISL_416480|2020-03-11

NL|Gelderland1|EPI_ISL_415461|2020-03-10

NL|Utrecht17|EPI_ISL_415526|2020-03-09

NL|Utrecht_16|EPI_ISL_414555|2020-03-08

BE|GHB-03021|EPI_ISL_407976|2020-02-03

NL|ZuidHolland27|EPI_ISL_415531|2020-03-09

NL|NA6|EPI_ISL_415495|2020-03-10

NL|Limburg7|EPI_ISL_415464|2020-03-09

BE|SN-03031|EPI_ISL_416469|2020-03-03

NL|NA23|EPI_ISL_415480|2020-03-10

NL|ZuidHolland_9|EPI_ISL_414445|2020-03-03

NL|Utrecht_1363628|EPI_ISL_413589|2020-03-01

BE|DB-03023|EPI_ISL_416470|2020-03-02

NL|NA17|EPI_ISL_415473|2020-03-10

NL|NA18|EPI_ISL_415474|2020-03-10

NL|NA28|EPI_ISL_415485|2020-03-10

GE|Tb-273|EPI_ISL_416479|2020-03-05

GE|Tb-673|EPI_ISL_416478|2020-03-14

IE|Limerick-19934|EPI_ISL_414586|2020-03-03

HU|mbl1|EPI_ISL_416426|2020-03-17

NL|Haarlem_1363688|EPI_ISL_413572|2020-03-01

NL|NoordBrabant_17|EPI_ISL_414457|2020-03-06

NL|ZuidHolland31|EPI_ISL_415535|2020-03-09

NL|NoordHolland3|EPI_ISL_415525|2020-03-09

NL|NA31|EPI_ISL_415488|2020-03-10

NL|NA30|EPI_ISL_415487|2020-03-10

NL|NA24|EPI_ISL_415481|2020-03-10

NL|ZuidHolland_17|EPI_ISL_414559|2020-03-07

NL|NA16|EPI_ISL_415472|2020-03-10

NL|Gelderland2|EPI_ISL_415462|2020-03-09

NL|Flevoland1|EPI_ISL_415460|2020-03-09

NL|NA1|EPI_ISL_415465|2020-03-10

NL|NA10|EPI_ISL_415466|2020-03-10

NL|NA13|EPI_ISL_415469|2020-03-10

GE|Tb|EPI_ISL_416482|2020-03-13

DK|SSI-09|EPI_ISL_416141|2020-03-03

NL|Utrecht_11|EPI_ISL_414443|2020-03-03

NL|Blaricum_1364780|EPI_ISL_413566|2020-03-02

NL|Zeewolde_1365080|EPI_ISL_413591|2020-03-02

BE|MTR-03026|EPI_ISL_416476|2020-03-02

BE|MTR-03021|EPI_ISL_416467|2020-03-02

NL|NA11|EPI_ISL_415467|2020-03-10

NL|NA26|EPI_ISL_415483|2020-03-10

NL|NA9|EPI_ISL_415498|2020-03-10

NL|NA8|EPI_ISL_415497|2020-03-10

NL|ZuidHolland_20|EPI_ISL_414562|2020-03-03

NL|ZuidHolland_15|EPI_ISL_414557|2020-03-08

NL|ZuidHolland_19|EPI_ISL_414561|2020-03-05

NL|NA12|EPI_ISL_415468|2020-03-10

NL|NoordBrabant55|EPI_ISL_415511|2020-03-09

NL|ZuidHolland26|EPI_ISL_415530|2020-03-09

NL|Utrecht_15|EPI_ISL_414554|2020-03-08

NL|NA25|EPI_ISL_415482|2020-03-10

NL|NA19|EPI_ISL_415475|2020-03-10

NL|NA15|EPI_ISL_415471|2020-03-10

BE|DBA-03032|EPI_ISL_416475|2020-03-03

BE|GMH-03022|EPI_ISL_416468|2020-03-02

NL|Gelderland3|EPI_ISL_415463|2020-03-09

NL|NA7|EPI_ISL_415496|2020-03-10

NL|NoordBrabant45|EPI_ISL_415502|2020-03-09

NL|ZuidHolland25|EPI_ISL_415529|2020-03-09

NL|ZuidHolland29|EPI_ISL_415533|2020-03-10

NL|Utrecht_13|EPI_ISL_414552|2020-03-07

NL|ZuidHolland_22|EPI_ISL_414564|2020-03-08

NL|ZuidHolland_10|EPI_ISL_414446|2020-02-26

NL|Rotterdam_1364740|EPI_ISL_413584|2020-03-02

DK|SSI-05|EPI_ISL_416140|2020-03-03

NL|ZuidHolland28|EPI_ISL_415532|2020-03-09

BE|DBD-03024|EPI_ISL_416471|2020-03-02

BE|UMF-03025|EPI_ISL_416472|2020-03-02

NL|ZuidHolland_13|EPI_ISL_414470|2020-03-06

NL|NA2|EPI_ISL_415476|2020-03-10

NL|Nootdorp_1364222|EPI_ISL_413579|2020-03-03

NL|Tilburg_1364286|EPI_ISL_413587|2020-03-03

IE|Limerick-19935|EPI_ISL_414587|2020-03-03

GE|Tb-477|EPI_ISL_415642|2020-03-10

GE|Tb-468|EPI_ISL_415643|2020-03-02

FI|FIN-455|EPI_ISL_414642|2020-03-08

FI|FIN0303C|EPI_ISL_413604|2020-03-03

GE|Tb-390|EPI_ISL_416477|2020-03-08

FI|FIN-313|EPI_ISL_414641|2020-03-05

FI|FIN0303A|EPI_ISL_413602|2020-03-03

FI|FIN-508|EPI_ISL_414643|2020-03-07

FI|FIN-266|EPI_ISL_414646|2020-03-04

KR|SNU01|EPI_ISL_411929|2020-01-NA

JP|DP0654|EPI_ISL_416613|2020-02-17

JP|DP0703|EPI_ISL_416619|2020-02-17

JP|DP0005|EPI_ISL_416565|2020-02-15

JP|DP0464|EPI_ISL_416603|2020-02-16

JP|DP0476|EPI_ISL_416604|2020-02-16

JP|DP0278|EPI_ISL_416587|2020-02-16

JP|DP0191|EPI_ISL_416582|2020-02-15

JP|TK20-31-3|EPI_ISL_413459|2020-02-20

JP|DP0588|EPI_ISL_416610|2020-02-17

JP|DP0357|EPI_ISL_416598|2020-02-16

JP|Hu_DP_Kng_19-027|EPI_ISL_412969|2020-02-10

KR|KCDC05|EPI_ISL_412869|2020-01-30

KR|KCDC24|EPI_ISL_412873|2020-02-06

JP|DP0037|EPI_ISL_416567|2020-02-15

JP|AII-004|EPI_ISL_407084|2020-01-25

KR|KCDC07|EPI_ISL_412871|2020-01-31

KR|KCDC06|EPI_ISL_412870|2020-01-30

JP|TY-WK-012|EPI_ISL_408665|2020-01-29

JP|TY-WK-521|EPI_ISL_408667|2020-01-31

JP|TY-WK-501|EPI_ISL_408666|2020-01-31

JP|DP0065|EPI_ISL_416570|2020-02-15

KR|KUMC01|EPI_ISL_413017|2020-02-06

KR|KCDC03|EPI_ISL_407193|2020-01-25

KR|KUMC02|EPI_ISL_413018|2020-02-06

JP|TKYE6182|EPI_ISL_414511|2020-01-00

JP|KY-V-029|EPI_ISL_408669|2020-01-29

JP|DP0481|EPI_ISL_416605|2020-02-16

JP|DP0287|EPI_ISL_416589|2020-02-16

JP|DP0457|EPI_ISL_416601|2020-02-16

JP|DP0200|EPI_ISL_416584|2020-02-15

JP|DP0361|EPI_ISL_416599|2020-02-16

JP|DP0027|EPI_ISL_416566|2020-02-15

JP|DP0190|EPI_ISL_416581|2020-02-15

JP|DP0158|EPI_ISL_416579|2020-02-15

JP|DP0236|EPI_ISL_416585|2020-02-15

JP|DP0724|EPI_ISL_416620|2020-02-17

JP|DP0880|EPI_ISL_416633|2020-02-17

KR|KCDC12|EPI_ISL_412872|2020-02-01

JP|DP0543|EPI_ISL_416607|2020-02-17

JP|DP0544|EPI_ISL_416608|2020-02-17

JP|DP0077|EPI_ISL_416571|2020-02-15

JP|DP0311|EPI_ISL_416593|2020-02-16

JP|DP0196|EPI_ISL_416583|2020-02-15

JP|DP0743|EPI_ISL_416621|2020-02-17

JP|DP0152|EPI_ISL_416578|2020-02-15

JP|DP0687|EPI_ISL_416614|2020-02-17

JP|DP0645|EPI_ISL_416612|2020-02-17

JP|DP0290|EPI_ISL_416591|2020-02-16

JP|DP0059|EPI_ISL_416569|2020-02-15

JP|DP0107|EPI_ISL_416574|2020-02-15

JP|DP0134|EPI_ISL_416577|2020-02-15

JP|DP0289|EPI_ISL_416590|2020-02-16

JP|DP0319|EPI_ISL_416594|2020-02-16

JP|DP0344|EPI_ISL_416596|2020-02-16

JP|DP0346|EPI_ISL_416597|2020-02-16

JP|DP0438|EPI_ISL_416600|2020-02-16

JP|DP0568|EPI_ISL_416609|2020-02-17

JP|DP0644|EPI_ISL_416611|2020-02-17

JP|DP0804|EPI_ISL_416631|2020-02-17

JP|DP0827|EPI_ISL_416632|2020-02-17

JP|DP0078|EPI_ISL_416572|2020-02-15

JP|DP0104|EPI_ISL_416573|2020-02-15

JP|DP0121|EPI_ISL_416575|2020-02-15

JP|DP0133|EPI_ISL_416576|2020-02-15

JP|DP0184|EPI_ISL_416580|2020-02-15

JP|DP0274|EPI_ISL_416586|2020-02-15

JP|DP0294|EPI_ISL_416592|2020-02-16

JP|DP0328|EPI_ISL_416595|2020-02-16

JP|DP0482|EPI_ISL_416606|2020-02-16

JP|DP0699|EPI_ISL_416617|2020-02-17

JP|DP0890|EPI_ISL_416634|2020-02-17

JP|Hu_DP_Kng_19-020|EPI_ISL_412968|2020-02-10

IN|1-27|EPI_ISL_413522|2020-01-27

NZ|20VR0189|EPI_ISL_416519|2020-03-02

AU|VIC08|EPI_ISL_416514|2020-03-15

VN|CM99|EPI_ISL_416429|2020-02-11

AU|VIC01|EPI_ISL_406844|2020-01-25

IN|1-31|EPI_ISL_413523|2020-01-31

AU|QLD02|EPI_ISL_407896|2020-01-30

AU|QLD03|EPI_ISL_410717|2020-02-05

AU|QLD04|EPI_ISL_410718|2020-02-05

AU|QLD01|EPI_ISL_407894|2020-01-28

AU|AU2|EPI_ISL_408976|2020-01-22

VN|VR03-38142|EPI_ISL_408668|2020-01-24

SG|8|EPI_ISL_410714|2020-02-03

AU|NSW01|EPI_ISL_407893|2020-01-24

SG|11|EPI_ISL_410719|2020-02-02

AU|NSW10|EPI_ISL_413596|2020-02-28

AU|NSW08|EPI_ISL_413594|2020-02-28

SG|7|EPI_ISL_410713|2020-01-27

SG|10|EPI_ISL_410716|2020-02-04

SG|9|EPI_ISL_410715|2020-02-04

AU|3|EPI_ISL_408977|2020-01-25

SG|3|EPI_ISL_407988|2020-02-01

SG|1|EPI_ISL_406973|2020-01-23

VN|39607|EPI_ISL_416428|2020-03-07

VN|CM295|EPI_ISL_416430|2020-03-06

VN|CM296|EPI_ISL_416431|2020-03-06

NZ|01|EPI_ISL_413490|2020-02-27

AU|NSW09|EPI_ISL_413595|2020-02-28

AU|NSW12|EPI_ISL_413598|2020-03-04

AU|NSW11|EPI_ISL_413597|2020-03-02

AU|NSW06|EPI_ISL_413213|2020-02-29

AU|NSW13|EPI_ISL_413599|2020-03-04

AU|NSW07|EPI_ISL_413214|2020-02-29

AU|NSW14|EPI_ISL_413600|2020-03-03

AU|NSW05|EPI_ISL_412975|2020-02-28

TW|CGMH-CGU-04|EPI_ISL_415742|2020-02-27

CN|P0028|EPI_ISL_413864|2020-02-05

CN_Jiangsu|IVDC-JS-001|EPI_ISL_408488|2020-01-19

TW|CGMH-CGU-05|EPI_ISL_415743|2020-02-27

CN_Wuhan|HBCDC-HB-05|EPI_ISL_412981|2020-01-18

TW|NTU03|EPI_ISL_413592|2020-02-29

CN_Guangzhou|GZMU0030|EPI_ISL_414684|2020-02-27

CN_Guangzhou|GZMU0030|EPI_ISL_414685|2020-02-27

CN_Guangzhou|GZMU0030|EPI_ISL_414686|2020-02-27

HK|VM20002907|EPI_ISL_414517|2020-02-25

HK|case85_VM20002868|EPI_ISL_414519|2020-02-24

CN_Jingzhou|HBCDC-HB-01|EPI_ISL_412459|2020-01-08

HK|case42_VM20002493|EPI_ISL_414527|2020-02-09

HK|case48_VM20002507|EPI_ISL_414528|2020-02-10

HK|VM20001988|EPI_ISL_412029|2020-01-30

HK|VM20002582|EPI_ISL_414569|2020-02-12

CN_Jiangsu|JS02|EPI_ISL_411952|2020-01-24

CN_Fujian|FJ13|EPI_ISL_411066|2020-01-22

CN_Foshan|20SF211|EPI_ISL_406536|2020-01-22

CN_Foshan|20SF210|EPI_ISL_406535|2020-01-22

TW|2|EPI_ISL_406031|2020-01-23

CN_Guangzhou|GZMU0031|EPI_ISL_414687|2020-02-25

CN_Guangzhou|GZMU0014|EPI_ISL_414692|2020-02-25

CN_Wuhan|IVDC-HB-envF13-21|EPI_ISL_408515|2020-01-01

CN_Jiangsu|JS01|EPI_ISL_411950|2020-01-23

CN_Wuhan|IPBCAMS-WH-01|EPI_ISL_402123|2019-12-24

CN_Zhejiang|WZ-01|EPI_ISL_404227|2020-01-16

CN|WF0003|EPI_ISL_413693|2020-01-NA

CN_Guangzhou|IQTC01|EPI_ISL_412966|2020-02-05

CN_Wuhan|WIV07|EPI_ISL_402130|2019-12-30

CN|WF0002|EPI_ISL_413692|2020-02-02

CN|WF0004|EPI_ISL_413694|2020-01-NA

CN_Wuhan|WH01|EPI_ISL_406798|2019-12-26

TW|NTU02|EPI_ISL_410218|2020-02-05

CN_Wuhan|IVDC-HB-05|EPI_ISL_402121|2019-12-30

CN_Shenzhen|SZTH-003|EPI_ISL_406594|2020-01-16

CN_Wuhan|IVDC-HB-04|EPI_ISL_402120|2020-01-01

HK|case78_VM20002849|EPI_ISL_414571|2020-02-22

CN_Wuhan|WIV02|EPI_ISL_402127|2019-12-30

HK|VB20026565|EPI_ISL_412030|2020-02-01

CN_Guangdong|20SF040|EPI_ISL_403937|2020-01-18

CN_Guangdong|20SF028|EPI_ISL_403936|2020-01-17

CN_Wuhan|HBCDC-HB-02|EPI_ISL_412898|2019-12-30

CN_Wuhan|HBCDC-HB-01|EPI_ISL_402132|2019-12-30

CN_Chongqing|IVDC-CQ-001|EPI_ISL_408481|2020-01-18

CN_Wuhan|IVDC-HB-envF13-20|EPI_ISL_408514|2020-01-01

CN_Wuhan|IPBCAMS-WH-05|EPI_ISL_403928|2020-01-01

CN_Wuhan|IPBCAMS-WH-03|EPI_ISL_403930|2019-12-30

CN_Anhui|SZ005|EPI_ISL_413485|2020-01-24

CN|WF0012|EPI_ISL_413697|2020-01-NA

CN|WF0026|EPI_ISL_413761|2020-02-NA

CN|WF0028|EPI_ISL_413791|2020-02-NA

HK|VM20001061|EPI_ISL_412028|2020-01-22

CN_Guangzhou|GZMU0042|EPI_ISL_414688|2020-02-25

CN_Shenzhen|HKU-SZ-005|EPI_ISL_405839|2020-01-11

CN|WF0029|EPI_ISL_413809|2020-02-NA

CN_Guangdong|20SF025|EPI_ISL_403935|2020-01-15

CN_Shenzhen|SZTH-002|EPI_ISL_406593|2020-01-13

CN_Shenzhen|HKU-SZ-002|EPI_ISL_406030|2020-01-10

CN|WF0021|EPI_ISL_413751|2020-02-NA

CN_Shandong|LY005|EPI_ISL_414938|2020-01-24

CN_Wuhan|HBCDC-HB-06|EPI_ISL_412982|2020-02-07

CN_Wuhan|HBCDC-HB-04|EPI_ISL_412980|2020-01-18

CN_Beijing|235|EPI_ISL_413521|2020-01-28

CN_Beijing|231|EPI_ISL_413519|2020-01-28

CN_Beijing|105|EPI_ISL_413518|2020-01-26

CN_Beijing|233|EPI_ISL_413520|2020-01-28

CN_Shandong|LY003|EPI_ISL_414936|2020-01-23

CN_Wuhan|HBCDC-HB-02|EPI_ISL_412978|2020-01-17

CN|WF0014|EPI_ISL_413711|2020-01-NA

CN|WF0019|EPI_ISL_413749|2020-02-NA

CN_Chongqing|YC01|EPI_ISL_408478|2020-01-21

CN_Shandong|LY008|EPI_ISL_414941|2020-01-30

CN_Fujian|FJ8|EPI_ISL_411060|2020-01-21

CN_Wuhan|WH04|EPI_ISL_406801|2020-01-05

CN_Yunnan|IVDC-YN-003|EPI_ISL_408480|2020-01-17

CN_Wuhan|HBCDC-HB-03|EPI_ISL_412979|2020-01-18

TW|3|EPI_ISL_411926|2020-01-24

CN|WF0020|EPI_ISL_413750|2020-02-NA

CN|WF0015|EPI_ISL_413729|2020-02-NA

CN|WF0001|EPI_ISL_413691|2020-02-02

CN_Guangzhou|GZMU0044|EPI_ISL_414689|2020-02-25

CN|WF0016|EPI_ISL_413746|2020-02-NA

CN|WF0018|EPI_ISL_413748|2020-02-NA

CN_Guangzhou|GZMU0048|EPI_ISL_414691|2020-02-25

CN_Shandong|LY007|EPI_ISL_414940|2020-01-25

CN|P0042|EPI_ISL_413854|2020-01-30

CN_Hangzhou|ZJU-08|EPI_ISL_416473|2020-01-26

CN_Hangzhou|ZJU-01|EPI_ISL_415709|2020-01-25

CN_Shenzhen|SZTH-001|EPI_ISL_406592|2020-01-13

CN_Guangzhou|IQTC02|EPI_ISL_412967|2020-01-29

CN_Shanghai|SH01|EPI_ISL_414510|2020-02-02

CN_Shenzhen|SZTH-004|EPI_ISL_406595|2020-01-16

CN_Hangzhou|ZJU-07|EPI_ISL_416425|2020-02-03

CN_Hangzhou|ZJU-02|EPI_ISL_416042|2020-01-25

CN_Hangzhou|ZJU-05|EPI_ISL_415711|2020-01-22

CN_Wuhan|WH-09|EPI_ISL_411957|2020-01-08

CN_Shandong|IVDC-SD-001|EPI_ISL_408482|2020-01-19

CN_Jiangxi|IVDC-JX-002|EPI_ISL_408486|2020-01-11

CN_Sichuan|IVDC-SC-001|EPI_ISL_408484|2020-01-15

CN_Hangzhou|ZJU-03|EPI_ISL_416044|2020-01-25

CN_Hangzhou|ZJU-06|EPI_ISL_416047|2020-02-02

CN_Hangzhou|ZJU-09|EPI_ISL_416474|2020-01-28

CN_Hangzhou|ZJU-04|EPI_ISL_416046|2020-01-24

US|NY1-PV08001|EPI_ISL_414476|2020-02-29

US|CA-CDPH-UC1|EPI_ISL_413557|2020-02-00

US|CA-PC101P|EPI_ISL_414648|2020-03-08

US|CA-CDPH-UC3|EPI_ISL_413559|2020-02-27

US|WA-UW21|EPI_ISL_414369|2020-03-05

US|WA-UW41|EPI_ISL_415606|2020-03-10

US|WA-UW19|EPI_ISL_414367|2020-03-05

US|WA-UW30|EPI_ISL_414617|2020-03-09

US|WA-UW48|EPI_ISL_415613|2020-03-10

US|WA-UW49|EPI_ISL_415614|2020-03-10

US|CA7|EPI_ISL_411954|2020-02-06

US|WA-UW31|EPI_ISL_414618|2020-03-08

US|WA-UW15|EPI_ISL_414363|2020-03-04

US|WA1|EPI_ISL_404895|2020-01-19

US|WA1-F6|EPI_ISL_407215|2020-01-25

US|WA1-A12|EPI_ISL_407214|2020-01-25

US|WA6-UW3|EPI_ISL_413457|2020-02-29

US|WA18-UW14|EPI_ISL_413653|2020-03-05

US|WA12-UW8|EPI_ISL_413563|2020-03-03

US|UPHL-03|EPI_ISL_415541|2020-03-09

US|UPHL-04|EPI_ISL_415542|2020-03-09

US|WA7-UW4|EPI_ISL_413458|2020-03-01

US|CA-CDPH-UC9|EPI_ISL_413928|2020-03-05

US|MN3-MDH3|EPI_ISL_414590|2020-03-09

US|WA-S11|EPI_ISL_416466|2020-03-03

US|WA-UW26|EPI_ISL_414595|2020-03-05

US|WA15-UW11|EPI_ISL_413650|2020-03-05

US|WA16-UW12|EPI_ISL_413651|2020-03-05

US|WA-UW18|EPI_ISL_414366|2020-03-05

US|WA-S10|EPI_ISL_416465|2020-02-29

US|WA-UW64|EPI_ISL_415592|2020-03-10

US|CA1|EPI_ISL_406034|2020-01-23

US|IL2|EPI_ISL_410045|2020-01-28

US|IL1|EPI_ISL_404253|2020-01-21

US|CruiseA-24|EPI_ISL_414483|2020-02-17

US|UPHL-02|EPI_ISL_415540|2020-03-09

US|CA-MG0987|EPI_ISL_416457|2020-03-18

US|UPHL-05|EPI_ISL_415543|2020-03-09

US|UPHL-06|EPI_ISL_415544|2020-03-09

US|WI1|EPI_ISL_408670|2020-01-31

US|CruiseA-23|EPI_ISL_414482|2020-02-18

US|CruiseA-21|EPI_ISL_414480|2020-02-21

US|TX1|EPI_ISL_411956|2020-02-11

US|MA1|EPI_ISL_409067|2020-01-29

US|CruiseA-26|EPI_ISL_414485|2020-02-24

US|CA3|EPI_ISL_408008|2020-01-29

US|CA4|EPI_ISL_408009|2020-01-29

US|AZ1|EPI_ISL_406223|2020-01-22

US|CruiseA-14|EPI_ISL_413619|2020-02-25

US|CruiseA-6|EPI_ISL_413611|2020-02-21

US|CruiseA-7|EPI_ISL_413612|2020-02-17

US|CA5|EPI_ISL_408010|2020-01-29

US|CruiseA-4|EPI_ISL_413609|2020-02-21

US|CruiseA-2|EPI_ISL_413607|2020-02-18

US|CruiseA-1|EPI_ISL_413606|2020-02-17

US|UPHL-01|EPI_ISL_415539|2020-03-09

US|CA2|EPI_ISL_406036|2020-01-22

US|CA8|EPI_ISL_411955|2020-02-10

US|CA9|EPI_ISL_412862|2020-02-23

US|CruiseA-17|EPI_ISL_413622|2020-02-24

US|CruiseA-12|EPI_ISL_413617|2020-02-20

US|CruiseA-11|EPI_ISL_413616|2020-02-17

US|CruiseA-10|EPI_ISL_413615|2020-02-17

US|CruiseA-8|EPI_ISL_413613|2020-02-17

US|CruiseA-25|EPI_ISL_414484|2020-02-17

US|UC-CDPH-UC11|EPI_ISL_413931|2020-03-05

US|CA-CDPH-UC2|EPI_ISL_413558|2020-02027

US|WA-S3|EPI_ISL_413560|2020-02-28

US|WA2|EPI_ISL_412970|2020-02-24

US|WA-S2|EPI_ISL_413456|2020-02-20

US|WA-S5|EPI_ISL_416460|2020-02-29

US|WA-S6|EPI_ISL_416461|2020-02-29

US|WA-UW32|EPI_ISL_414619|2020-03-07

US|WA-UW60|EPI_ISL_415625|2020-03-10

US|WA-UW22|EPI_ISL_414591|2020-03-06

US|WA-UW69|EPI_ISL_415597|2020-03-10

US|WA-UW73|EPI_ISL_415601|2020-03-10

US|WA-UW24|EPI_ISL_414593|2020-03-05

US|WA-UW74|EPI_ISL_415602|2020-03-10

US|WA-UW44|EPI_ISL_415609|2020-03-10

US|WA-UW57|EPI_ISL_415622|2020-03-10

US|MN2-MDH2|EPI_ISL_414589|2020-03-07

US|WA-UW28|EPI_ISL_414597|2020-03-04

US|WA-UW62|EPI_ISL_415627|2020-03-10

US|WA-UW43|EPI_ISL_415608|2020-03-10

US|WA-UW63|EPI_ISL_415591|2020-03-09

US|WA-UW46|EPI_ISL_415611|2020-03-10

US|WA-UW33|EPI_ISL_414620|2020-03-08

US|WA-UW59|EPI_ISL_415624|2020-03-10

US|WA-UW67|EPI_ISL_415595|2020-03-10

US|WA8-UW5|EPI_ISL_413486|2020-03-01

US|WA-UW56|EPI_ISL_415621|2020-03-10

US|WA-UW75|EPI_ISL_415603|2020-03-10

US|WA-UW76|EPI_ISL_415604|2020-03-10

US|WA-UW50|EPI_ISL_415615|2020-03-10

US|WA-UW66|EPI_ISL_415594|2020-03-10

US|WA-UW68|EPI_ISL_415596|2020-03-10

US|WA-UW70|EPI_ISL_415598|2020-03-10

US|WA-UW55|EPI_ISL_415620|2020-03-10

US|WA-UW23|EPI_ISL_414592|2020-03-06

US|WA-UW20|EPI_ISL_414368|2020-03-05

BR|SPBR-14|EPI_ISL_416036|2020-03-05

MX|CDMX-InDRE_01|EPI_ISL_412972|2020-02-27

BR|SPBR-12|EPI_ISL_416034|2020-03-04

BR|SPBR-07|EPI_ISL_416028|2020-03-03

BR|SPBR-08|EPI_ISL_416029|2020-03-04

BR|SPBR-13|EPI_ISL_416035|2020-03-05

CL|Santiago-2|EPI_ISL_414580|2020-03-05

CL|Talca-1|EPI_ISL_414577|2020-03-02

CL|Talca-2|EPI_ISL_414578|2020-03-04

CL|Santiago-1|EPI_ISL_414579|2020-03-03

CL|Santiago_op2d1|EPI_ISL_415658|2020-03-06

CL|Santiago_op3d1|EPI_ISL_415660|2020-03-07

CL|Santiago_op4d1|EPI_ISL_415661|2020-03-08

BR|SPBR-09|EPI_ISL_416031|2020-03-04

BR|314|EPI_ISL_414045|2020-03-04

BR|SPBR-10|EPI_ISL_416032|2020-03-05

BR|SPBR-11|EPI_ISL_416033|2020-03-03

RS- MT510747

SP- MT292569

PK- MT240479
